# Incorporating climate velocity into the design of climate-smart networks of protected areas

**DOI:** 10.1101/2020.06.08.139519

**Authors:** Nur Arafeh-Dalmau, Isaac Brito-Morales, David S. Schoeman, Hugh P. Possingham, Carissa J. Klein, Anthony J. Richardson

**Affiliations:** Centre for Biodiversity and Conservation Science, School of Biological Sciences, The University of Queensland, St Lucia, Queensland, Australia; School of Earth and Environmental Sciences, The University of Queensland, St Lucia, Queensland, Australia; Commonwealth Scientific and Industrial Research Organisation (CSIRO) Oceans and Atmosphere, BioSciences Precinct (QBP), St Lucia, Queensland, Australia; Global-Change Ecology Research Group, School of Science and Engineering, University of the Sunshine Coast, Maroochydore, Queensland, Australia; Centre for African Conservation Ecology, Department of Zoology, Nelson Mandela University, Port Elizabeth, South Africa; The Nature Conservancy, Arlington, VA 22203–1606, USA; Centre for Applications in Natural Resource Mathematics, School of Mathematics and Physics, The University of Queensland, St Lucia, Queensland, Australia

**Keywords:** climate change, reserves, conservation planning, marine protected areas, climate adaptation, post-2020 conservation targets

## Abstract

Climate change is redistributing terrestrial and marine biodiversity and altering fundamental ecological interactions. To adequately conserve biodiversity and promote its long-term persistence, protected areas should account for the ecological implications of species’ redistribution. Data paucity across many systems means that achieving this goal requires generic metrics that represent likely responses of multiple taxa to climate change. Climate velocity is one such metric, reflecting potential species’ range shifts at a generic level. Here, we explore four approaches to incorporating climate velocity metrics into the design of protected areas using the Mediterranean Sea as an illustrative example. Our methods are designed to meet two climate-smart planning objectives: 1) protect climate refugia by selecting slow-moving climate velocity areas, and 2) maintain the capacity of ecological systems to adapt by representing a suite of climate-velocity trajectory classes. We found that incorporating climate velocity as a cost measure in Marxan is the best approach for selecting slower-moving areas, which are good indicators of climate refugia. However, this approach fails to accommodate socio-economic cost data, and is probably impractical. Incorporating climate velocity as a boundary or as a feature provides both selection of slower-moving areas and solutions with lower socio-economic cost. Finally, we were able to design cost-effective networks of protected areas representing a suite of climate-velocity trajectories classes, which have the potential to help species adapt to a changing climate. This work presents simple and practical ways of including climate velocity in conservation plans on land and in the ocean to achieve the key climate-smart objectives of protecting climate refugia and enhancing ecological resilience.

## 1. INTRODUCTION

Protected areas are the cornerstone of most conservation strategies as they are one of the most effective tools for protecting biodiversity (Edgar et al., 2014; Gray et al., 2016; Lubchenco et al., 2003; Watson et al., 2014). Spatial conservation prioritization is among the most common approaches used to support the design of networks of protected areas that achieve conservation and socioeconomic goals (Moilanen et al., 2009). However, although data on current and future ecological processes are critical when identifying protected areas, climate change and its impacts are rarely incorporated into spatial conservation prioritizations (Álvarez-Romero et al., 2018; Jones et al., 2016; Reside et al., 2018; Tittensor et al., 2019; Wilson et al., 2020).

Protected areas help species adapt to a changing climate (Duarte et al., 2020; Roberts et al., 2017; Webster et al., 2017). As climate warms, species shift their biogeographic distributions to track their preferred thermal niches (Pecl et al., 2017). Implications of this redistribution for biodiversity are twofold: encouraging the movement of species beyond the boundaries of existing protected areas (Araújo et al., 2004; Bates et al., 2019); and simultaneously altering fundamental ecological interactions and creating new ones (Pecl et al., 2017). Accounting for these processes when designing protected areas will promote the long-term persistence of biodiversity (Fredston-Hermann et al., 2018).

Where climate change is incorporated into spatial plans, this is usually achieved by prioritizing “future habitats” or “representing climate refugia” identified by species distribution models projected into the future based on climate-change scenarios (Jones et al., 2016; Lawler et al., 2020; Morelli et al., 2016). Despite significant advances, prohibitive costs of collecting biotic data, especially from the ocean, result in data deficiencies that limit the development of predictive models for the majority of species (Sequeira et al., 2018). Moreover, there are questions about how transferable these models are across both space and time (Robinson et al., 2011; Yates et al., 2018). A potential solution would be to use a suite of generic metrics, such as climate velocity, that describe likely responses of multiple taxa to climate change (Brito-Morales et al., 2018).

Climate velocity describes the speed and direction that a species at a given point in space would need to move to remain within its climatic niche (Loarie et al., 2009). This metric has been used to inform potential species distribution shifts, providing information about where biodiversity might be rearranged under a changing climate (Burrows et al., 2011; Hiddink et al., 2012; Molinos et al., 2016). The few examples that have used climate velocity in spatial prioritization have focused on identifying terrestrial climate refugia, and do not consider other important factors when designing protected areas (Carroll et al., 2017; Haight & Hammill, 2020). For example, Caroll et al. (2017) compared the performance of six approaches, including climate velocity, to identify priority refugia for protection but did not represent biodiversity (at the species or habitat level) or incorporate socioeconomic costs. Similarly, Haight & Hammill (2020) used climate velocity metrics as a cost in prioritizing the protection of climate refugia for a range of terrestrial species, but also overlooked socioeconomic costs. To advance the inclusion of climate velocity into the framework for climate-smart protected areas, we need approaches that include current best-practice protected-area design principles such as adequate representation of biodiversity and minimizing socio-economic costs.

Climate-smart protected areas conform to general design guidelines and conceptual frameworks that account for climate-change adaptation objectives (Hansen et al., 2010; Stein et al., 2014; West et al., 2017; Wilson et al., 2020). The two most important of these objectives are: 1) protecting climate refugia (Keppel et al., 2012), usually defined as areas where biodiversity retreats to, persists in, or can potentially expand from when climate changes; and 2) protecting a range of areas where future ecosystems could be altered in different ways due to a changing climate (Stein et al., 2014; Tittensor et al., 2019; Walsworth et al., 2019; Wilson et al., 2020). However, no general framework yet exists to provide practical advice on how to meet these objectives at a community or biogeographic level (Fredston-Hermann et al., 2018). Given that changes in species’ interactions have been a cause of extirpations and population declines during previous periods of climate change (Cahill et al., 2013), prioritizing the protection of areas of slow climate velocity, which are good indicators of climate refugia (Sandel et al., 2011), could contribute significantly to achieving climate-smart status. This has the advantage of simultaneously selecting for areas within which species are likely to remain. So, selecting areas with slow-moving climate velocities not only seeks to conserve climate refugia, but also tends to create protected areas that might retain their biodiversity under climate change.

An extension of climate velocity – called climate-velocity trajectory classes – could also provide valuable indicators for climate-smart protected area design. These describe spatial pathways along which a given thermal niche moves through time, suggesting how climate-migrant taxa are likely to move as climate changes (Burrows et al., 2014). Classes of climate-velocity trajectories could highlight areas where changes in species interactions are more likely (Burrows et al., 2014). For example, climate sources might represent areas with fast rates of emigration that receive few or no climate migrants, leaving empty niches. Climate sinks might be disconnected from cooler areas, resulting in potential extirpations replaced by new migrants. Climate corridors are areas where climates converge and might accelerate novel species interactions due to rapid emigration and immigration. In areas with slow-moving or static trajectories, there might be few changes in species distribution, representing potential climate refugia (Fogarty et al., 2017). How ecological processes (such us adaptation, extinction, or competition) will be altered as a result of species redistribution is unknown, but classes of climate trajectories suggest areas where future ecosystems could be altered in different ways (Burrows et al., 2014). Given that species have adapted to previous climate change throughout their evolutionary history (Davis & Shaw, 2001; Sandel et al., 2011), protecting a range of climate-trajectory classes could enhance the capacity of ecological systems to adapt to a changing climate (Brito-Morales et al., 2018).

Here, we assess several methods of incorporating climate velocity metrics into the design of climate-smart systems of protected areas. Although we illustrate the methods using the Mediterranean Sea as a case study, the methods are equally applicable to both terrestrial and marine systems. We address two climate-smart planning objectives: 1) protect climate refugia by maximising the selection of areas of slow climate velocity; and 2) maintain the capacity of ecological systems to adapt to climate change by ensuring representation of all climate-velocity trajectory classes. We evaluate the resulting protected-area networks in terms of their median climate velocity and socioeconomic cost. Our aim is not to suggest a network of protected areas that could be implemented in the Mediterranean Sea, but rather to develop, evaluate and compare approaches using climate velocity metrics for incorporation into the climate-smart protected area toolkit.

## 2. MATERIALS AND METHODS

### 2.1 Case study

The Mediterranean Sea is considered a climate change “hot-spot” (Giorgi, 2006), warming two-to-three times faster than the ocean as a whole (Vargas-Yáñez et al., 2008). This semi-enclosed basin is home to a high proportion of endemic species, some of which have limited abilities to track their preferred thermal conditions because of geographic obstructions. We divide this planning domain into 4,649 square planning units of 0.25° (coastal squares were smaller as they were clipped to the coastline).

### 2.2 Climate velocity and climate-velocity trajectories classes

We calculated local climate velocity (after Burrows et al. (2011)) and climate-velocity trajectories (after Burrows et al. (2014)) for the period 2020-2100 using the VoCC R package (García Molinos et al., 2019) and sea surface temperature (SST) projections from the MPI-ESM1-2-HR model at 0.5° spatial resolution (Figure 1a-b). We used an intermediate climate scenario generated under one of the IPCC Shared Socio-Economic Pathways (SSPs), SSP2(4.5), which represents intermediate challenges for mitigation with radiative forcing levels equivalent to RCP4.5 (radiative forcing is stabilized at ∼4.5 W m^-2^ by 2100). We did not use multiple future models and scenarios as is best practice for climate change research (Harris et al., 2014; Lotze et al., 2019), as our aim here is not to provide a comprehensive assessment of climate-change risks in the region, but to illustrate the strengths and weaknesses of different approaches to climate-smart protected area design.

**FIGURE 1.**
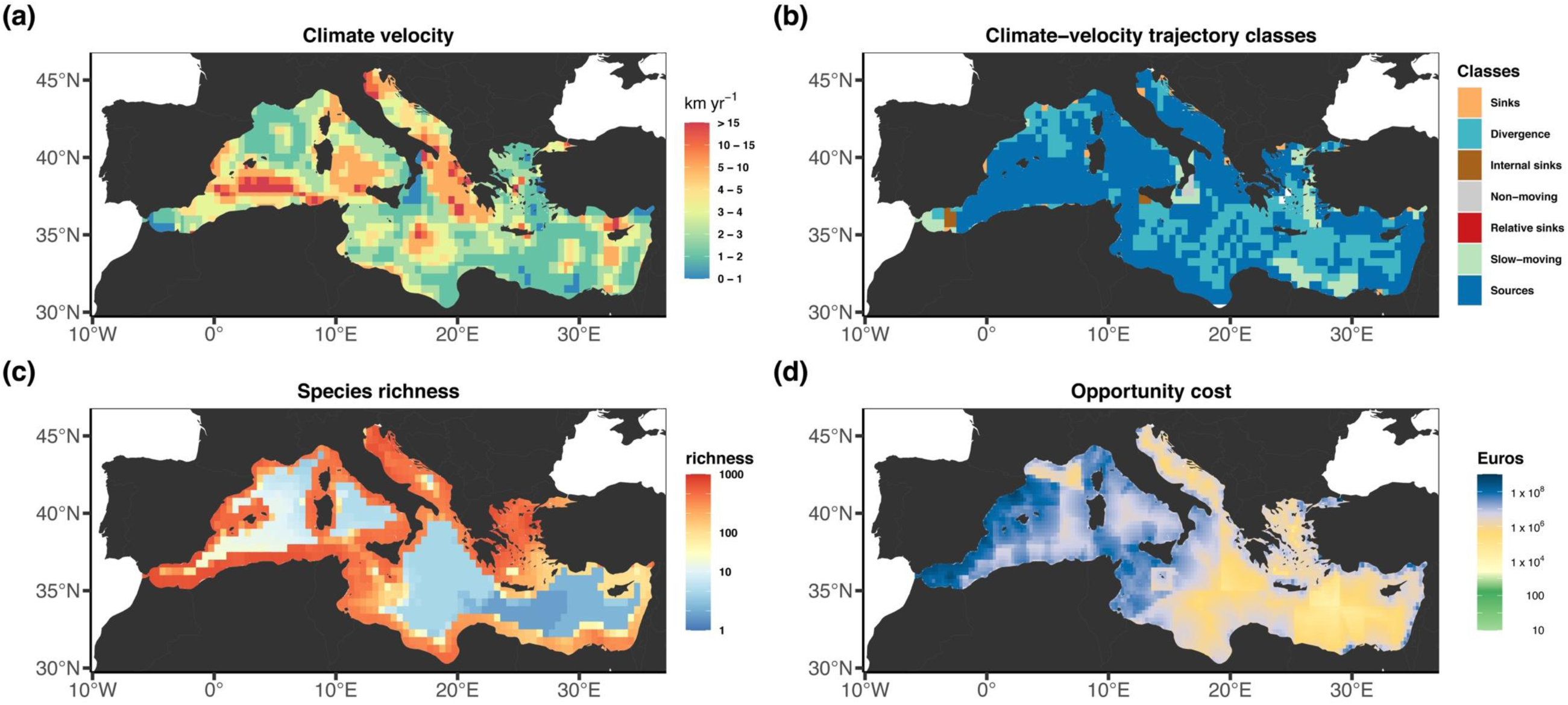
Maps of the study area quantifying the (a) climate velocity and (b) climate-velocity trajectory classes for the MPI-ESM1-2-HR Earth-system model under SSP2(4.5); (c) species richness and (d) the opportunity cost (Euros).

Climate-velocity trajectories were constructed for the same period as for climate velocity by forward iteration of isotherms located at the centre of each 0.5° cell, after which each cell was allocated to one of six different classes following Burrows et al. (2014): non-moving, slow-moving, internal sinks, boundary sinks, sources, relative sinks. To match the spatial resolution of climate velocity and climate-velocity trajectory classes with the scale of the planning units (0.25°), we disaggregated each cell into four cells of equal value and size.

### 2.3 Data for biodiversity features

We represented the biodiversity of our study region using AquaMaps data (Kaschner, 2019), which estimates the probability of occurrence (0-1) for each species at 0.5° spatial resolution derived from occurrence observations and environmental-niche models (based on depth, temperature, salinity, and oxygen). We used a threshold probability of 0.5 to compile range maps for our analysis (after (Klein et al., 2015). This yielded a total of 1,471 species distribution maps for the planning domain (Figure 1c and Table S1), which were used as conservation features to reflect the spatial representation of each species in each planning unit.

### 2.4 Opportunity costs

We used a surrogate cost layer for the Mediterranean Sea from Scenario 6 in Mazor et al. (2014). This represents the combined spatial opportunity cost in monetary value (Euros) for commercial (small- and large-scale) fishing, non-commercial (recreational and subsistence) fishing, and aquaculture activities (Figure 1d). More details are available in the supporting information. We aggregated the combined cost layer from 10 km^2^ to 0.25°grid cells to match our planning-unit size by applying a sum of the original cost values weighted by the proportion of each original cell in each planning unit.

### 2.5 Spatial Prioritization Approach

We used the spatial prioritization software Marxan to identify conservation priority areas to meet two climate-smart planning objectives (Table 1). Marxan uses the simulated annealing algorithm (Possingham et al., 2000) to find near-optimal solutions that meet a set of quantitative conservation targets, while minimizing the total perimeter of the reserve system and its total “cost”. The boundary-length modifier is a parameter in Marxan that allows users to control the compactness of the reserve system by adjusting the cost of the perimeter relative to the total opportunity cost.

**TABLE 1.**
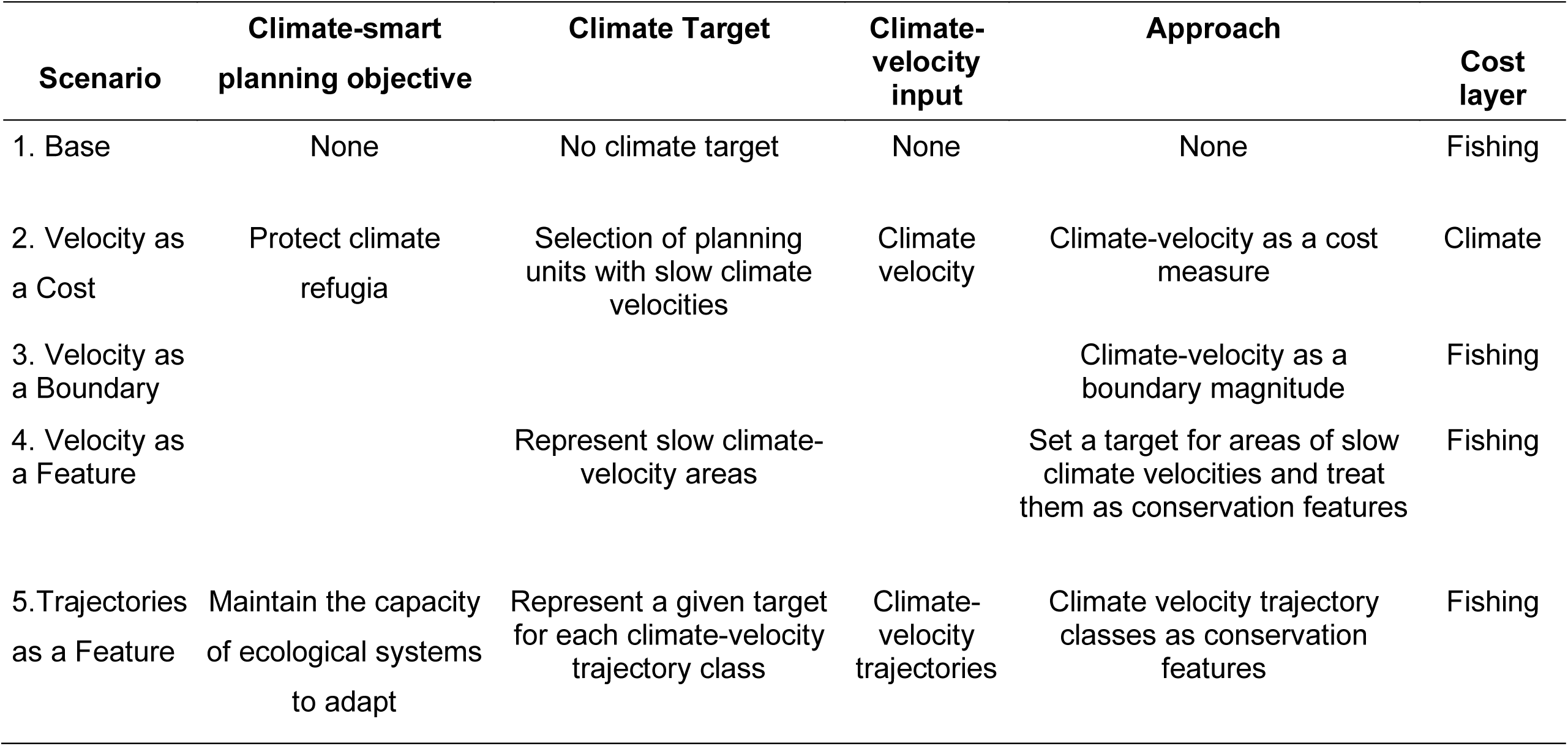
Spatial prioritization approaches (scenarios) based on climate smart-planning objectives, opportunity cost, and climate-change metrics.

### 2.6 Planning Scenarios

To plan for climate-smart protected areas, we constructed five scenarios that varied in their use and treatment of climate-velocity and opportunity-cost data (Table 1). All five scenarios represented 20% of each species’ distribution for a minimum cost (although cost varied by scenario). Scenario 1 – “Base”, did not attempt to meet any climate-smart planning objectives but included the opportunity cost. In the second to the fourth scenarios, we prioritized climate refugia based on climate velocity. In Scenario 2 – “Velocity as a Cost”, we followed Haight & Hammill (2020) by incorporating climate velocity as the cost layer (to minimize the selection of planning units with high climate velocities) in place of opportunity cost data. In Scenario 3 – “Velocity as a Boundary”, we used the magnitude of climate velocity as a boundary length to prioritize the selection of slow-velocity planning units while minimizing the opportunity cost (see next section, Figure S3). In Scenario 4 – “Velocity as a Feature”, we set a target to represent 30% of the slowest-moving planning units (defined as the first percentile of climate velocities). Finally, Scenario 5 – “Trajectories as a Feature”, we prioritized the representation of 20% of each climate-velocity trajectory class with the aim of maintaining the capacity of ecological systems to adapt.

For each scenario, we ran Marxan 100 times with 10 million iterations. We used a tailored approach to calibrate the number of iterations, the species penalty factor, and the boundary-length modifier (see supporting information, Figure S1-S2, Table S2, and http://doi.org/10.5281/zenodo.3876041 for the code). Finally, we also calibrated the representation target assigned to the slowest-moving planning units for the Velocity as a Feature scenario to find the optimal trade-off between cost, fragmentation, and area of the solution (see supporting information, Figure S4).

### 2.7 Climate velocity as a boundary

For the Velocity as a Boundary scenario, we developed an approach that modified the Marxan objective function to favour both the selection and the spatial aggregation of planning units with slow climate velocities. We identified the magnitude of climate velocity in each planning unit, then assigned these values as the boundary lengths of that unit. When planning units shared a boundary, we averaged the magnitudes of climate velocities of the respective neighbours. We could thus control the importance of climate velocity relative to the conservation targets and costs of the prioritization approach using the boundary-length modifier. A zero value for the boundary-length modifier means that there is no preference for selecting areas of lower or higher climate velocity, with an increasing value resulting in a greater emphasis on climate velocity in the selection of planning units to be included in the protected-area network. When applying non-zero values for the boundary-length modifier, Marxan will seek to minimize the sum of shared climate-velocity magnitudes. With these modifications, Marxan solves the conservation-planning problem of minimizing the sum of the total cost of selected sites, as well as the sum of the averaged climate-velocity magnitudes between shared boundaries across selected sites, subject to meeting the target for every biodiversity feature. Thus, a higher boundary-length modifier value will favour the selection and aggregation of lower velocity areas (Figure S3).

### 2.8 Comparing efficiency and spatial similarity among approaches

We evaluated the trade-offs among the different approaches used to prioritize climate refugia (Scenarios 2-4) and the Base scenario by assessing their efficiency in minimizing the median climate velocity and the total opportunity cost of selected planning units from 100 spatially different solutions to the problem. We also evaluated the spatial similarity (overlap) of the selection frequencies of the planning units among different scenarios. The selection frequency is a good indicator of the irreplaceability of planning units (Stewart & Possingham, 2005), and is commonly used to identify priority areas for conservation. We estimated Cohen’s kappa coefficient, a pairwise statistic (Goto & Watanabe, 2012) that indicates how much the selection frequencies of the planning units overlap among scenarios. Kappa ranges -1 to +1, where -1 indicates complete disagreement, 0 indicates overlap due to chance, and +1 indicates complete agreement (Landis & Koch, 1977). We followed the approach of (Ruiz-Frau et al., 2015) and classified planning-unit selection frequency into five classes (0, <25%, 25-50%, 50-75%, and >75%). Finally, we evaluated trade-offs for the Trajectories as a Feature scenario by assessing the efficiency of minimizing the total opportunity cost while meeting representation targets for each climate-velocity trajectory class compared with the Base scenario.

## 3. RESULTS & DISCUSSION

### 3.1 Protect climate refugia

The Base scenario was the best at minimizing the opportunity cost while meeting representation targets for all species, but it did a poor job at selecting slow-moving planning units, as it does not attempt to protect climate refugia (Figure 3). Velocity as a Cost was the most efficient and flexible scenario for minimizing the selection of slow-moving planning units and thus protecting climate refugia (Table 2), achieving the slowest median climate-velocity (1.675 ± 0.006 km yr^-1^) compared to the Base scenario (2.838 ± 0.022 km yr^-1^) (Figure 2a-b, Figure 3a). It also outperformed scenarios using Velocity as a Boundary (2.319 ± 0.035 km yr^-1^) and Velocity as a Feature (2.625 ± 0.013 km yr^-1^) (Figure 2c-d, and Figure 3a). However, because Velocity as a Cost did not include opportunity cost, it was least cost-effective (Table 2), increasing the total cost almost tenfold (€30,891.57 ± 266.14 million) relative to the Base scenario (€3,187.25 ± 10.81 million). In comparison, the Velocity as a Boundary (€4,748.98 ± 56.14 million) and Velocity as a Feature scenarios (3,176.51 ± 10.16 million) both had only a small increase in the total opportunity cost (Figure 3b).

**TABLE 2.** Assessment of the performance of three approaches developed to protect climate refugia. Flexibility is the freedom with which slow-moving planning units can be selected.

| Scenarios | Selecting<br>slow-moving | Cost-<br>efficiency | Flexibility | Recommendations |
| --- | --- | --- | --- | --- |
| Velocity as<br>a Cost | Best | Worst | Best | Identifies the best places to protect biodiversity in slow-moving areas. Not recommend, as it fails to minimize socio-economic cost. Flexible because selection of slowest planning units does not constrain the solution to a set of planning units or their spatial aggregation. |
| Velocity as<br>a Boundary | Intermediate | Intermediate | Intermediate | Recommended approach because it is best at finding cost-efficient solutions while selecting slow-moving planning units. Requires calibration of boundary-length modifier to choose appropriate trade-off when minimizing cost, compacting the solutions, and selecting slower velocity areas (see supporting information, Figure S3). Has intermediate flexibility because selection of slow-moving areas is constrained by spatial aggregation of planning units. |
| Velocity as<br>a Feature | Worst | Best | Worst | Requires a trade-off analysis to determine optimal: 1) threshold to classify slowest-moving planning units as a feature (i.e., between 5 to 25% of the slowest moving) that minimize median velocities without substantial increase in cost or area of Marxan solutions; and 2) representation targets for slow-moving features (see supporting information, Figure S4). |

**FIGURE 2.**
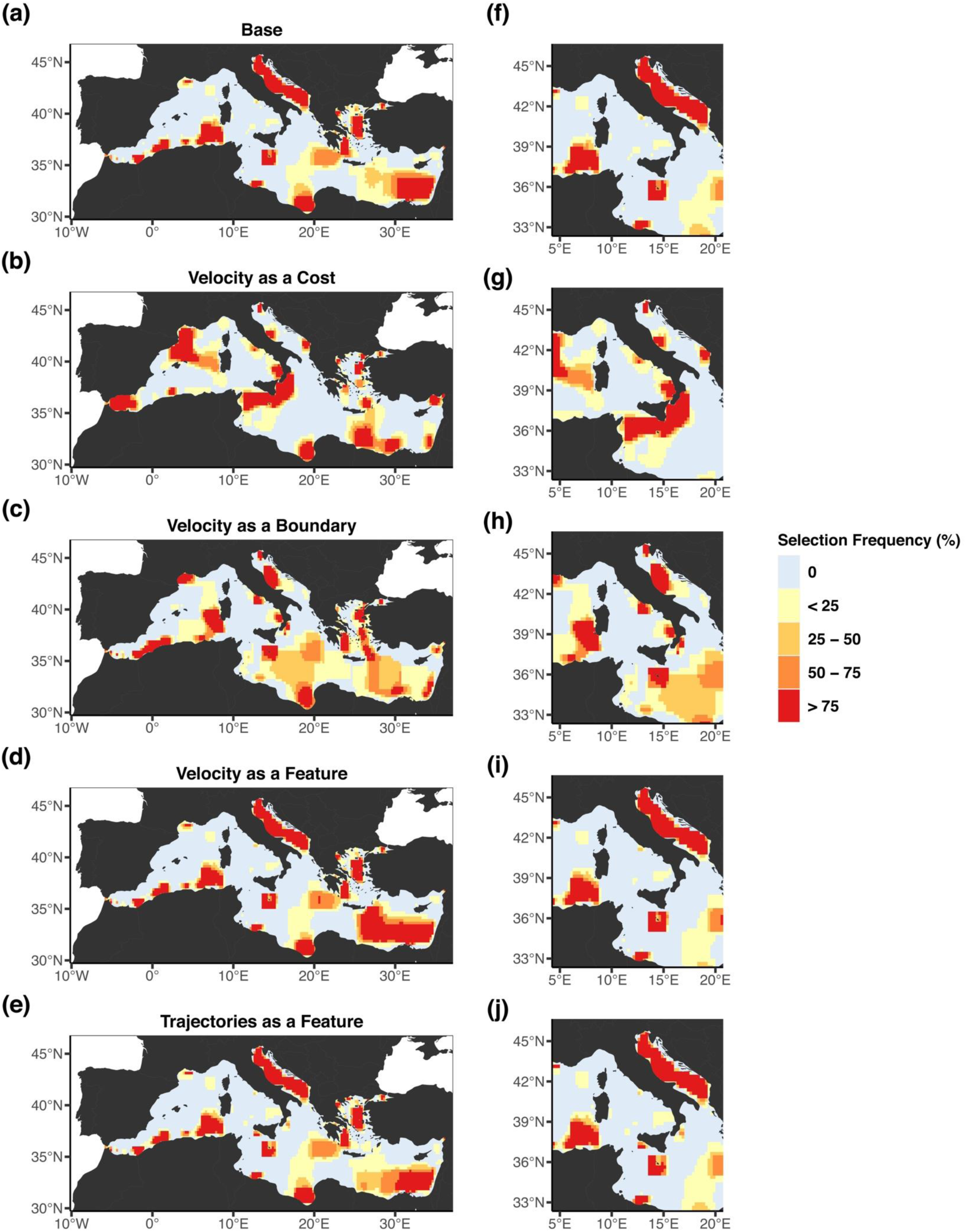
Selection frequency (the number of times a planning unit is selected in 100 Marxan solutions) for the Mediterranean Sea: (a), (f) Base scenario; (b), (g) Velocity as a Cost; (c), (h) Velocity as a Boundary; (d), (i) Velocity as a Feature; and (e), (j) Trajectories as a Feature. For each scenario, we present the full planning domain together with a magnified representation of the west and south coast of Italy and the Adriatic Sea.

**FIGURE 3.**
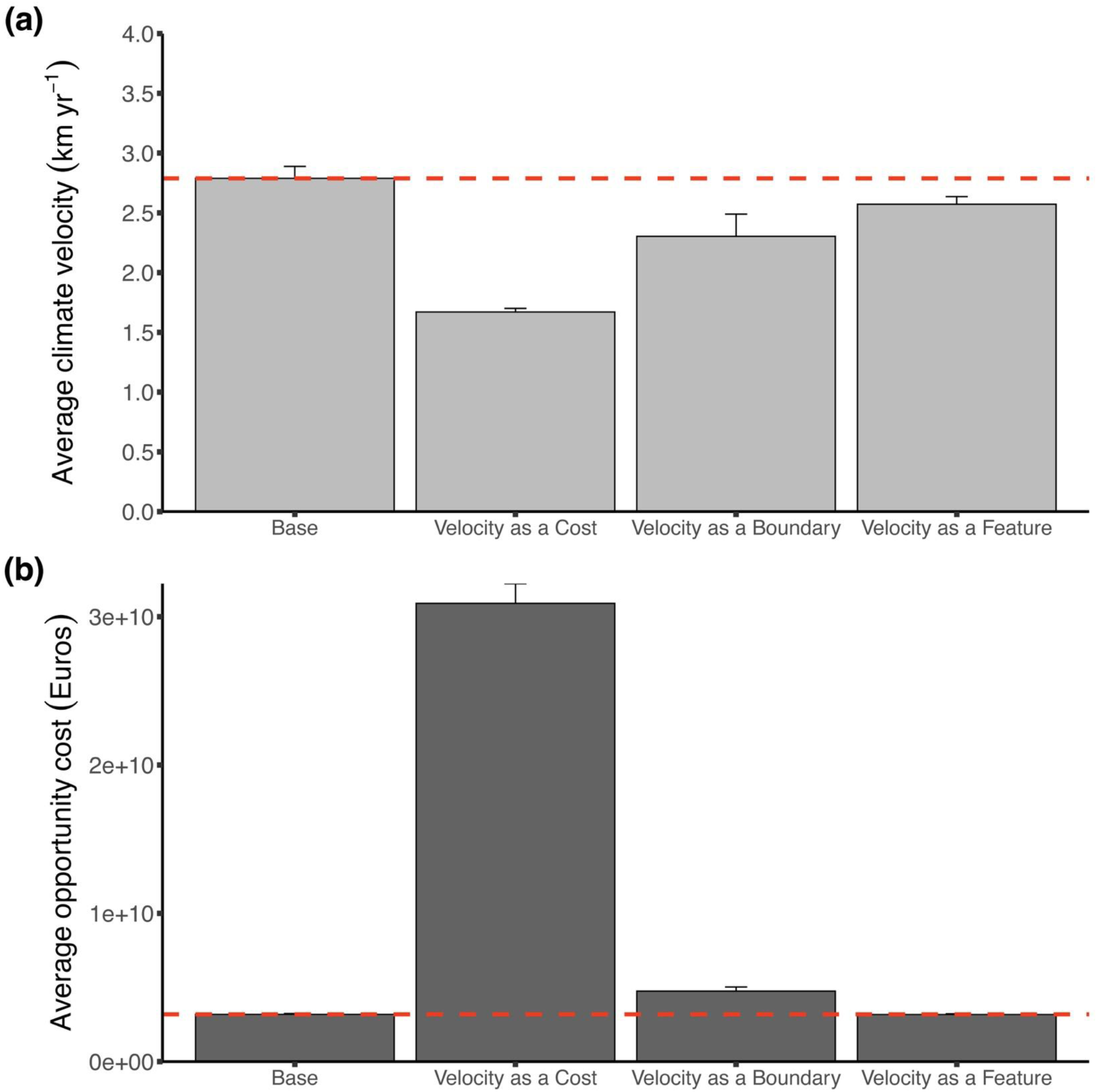
(a) Median climate velocity and (b) opportunity cost, Euros (n = 100 ± s.d) among Base scenario and scenarios designed to protect refugia (Velocity as a Cost, Velocity as a Boundary, Velocity as a Feature) for 100 solutions generated in Marxan. The horizontal dashed line indicates the average value for the base scenario.

A fundamental principle in conservation planning is to identify cost-effective priority areas that meet conservation targets, while minimizing potential human conflicts arising from the implementation of protected areas (Arafeh-Dalmau et al., 2017; Baker-Médard et al., 2019; Ban & Klein, 2009). As expected, networks of protected areas that used climate velocity as a cost (Figure 2b) yield inefficient economic solutions, and thus economically and politically non-viable climate-smart conservation plans. Both the Velocity as a Boundary and Velocity as a Feature scenarios appear to be good options; they minimized the selection of slow-moving planning units (Figure 2c-d), without dramatically increasing the opportunity cost.

While the Base scenario had moderate spatial overlap with Velocity as a Boundary and high overlap with Velocity as a Feature, the Velocity as a Cost scenario exhibited little overlap with these (Figure 2, Table 3). These results reiterate the principle that the opportunity cost controls the spatial configuration of solutions (Ban & Klein, 2009; Mazor et al., 2014), and also highlights the need to use approaches that meet conservation objectives while considering both socio-economic data and potential climate impacts.

**TABLE 3.** Spatial similarity matrix for prioritization approaches. Values represent Cohen’s kappa coefficient that ranges -1 to +1, where -1 indicates complete disagreement, 0 represents overlap due to chance, and +1 represents complete agreement.

| Scenarios | Base | Velocity as a Cost | Velocity as a Boundary | Velocity as a Feature |
| --- | --- | --- | --- | --- |
| Base | - |  |  |  |
| Velocity as Cost | 0.08 | - |  |  |
| Velocity as Boundary | 0.30 | 0.16 | - |  |
| Velocity as a Feature | 0.75 | 0.10 | 0.28 | - |

Velocity as a Boundary was the most cost-effective scenario for minimizing the selection of slow-moving areas, performing better than Velocity as a Feature (Figure 3, Table 2). However, when we calibrated Velocity as a Feature, we found that an increase in the representation target (from 40 to 70%) improved the selection of slow-moving areas, with little change in the cost, although it substantially increased the amount of area required to meet the targets (Figure S4), an undesired trade-off (Stewart & Possingham, 2005). The performance of this approach could be improved if we used a lower threshold for selecting slower-moving planning units to be represented as a feature (e.g., the lowest decile). Therefore, we recommend conducting a trade-off and calibration analysis (Table 2). An advantage of using Velocity as a Boundary is that it is a more flexible approach for selecting slow-moving areas because it considers all planning units, rather than being limited to a subset classified as “slow-moving”. We therefore recommend Velocity as a Boundary because it will always find solutions that minimize the overall magnitude of climate velocity without the need for further manipulation (Table 2). However, this comes with a slight increase in complexity of implementation relative to Velocity as a Feature.

Climate-smart conservation has started to prioritize retention of habitats within protected areas (Maxwell et al., 2019), an objective achieved by conserving slow-moving areas. Moreover, systems of protected areas that retain species within their boundaries should also better maintain the functioning of ecosystem (Fredston-Hermann et al., 2018) because species interactions experience least change when turnover of biodiversity is minimized (Burrows et al., 2014; Cahill et al., 2013; Molinos et al., 2016). These regions might be climate refugia (Keppel et al., 2012), which are areas characterized by either little environmental change or by a high diversity of environmental conditions, implying that resident species would not need to move far in the future to find an area with suitable conditions (Sandel et al., 2011).

### 3.2 Maintaining the capacity of ecosystems to adapt

We found that in the Trajectories as a Feature scenario, protecting 20% of each climate-velocity trajectory class (Figure 2E) came with little increase in the opportunity cost (€3,390.23 ± 9.06 million) compared to the Base scenario (€3,187.25 ± 10.81 million). Yet the Base scenario represented only three of the six climate-velocity trajectory classes (boundary sinks: 15.77 ± 2.97%, slow moving: 11.14 ± 3.42%, and sources: 20.34 ± 0.56%), and only sources met the representation target. Interestingly, of the 100 Marxan solutions, none represented non-moving trajectory classes or relative sinks (0%), and only one solution captured internal sinks (1.70%). We reiterate the importance of including representation targets for trajectory classes to ensure each is represented in a system of protected areas. Because some trajectory classes are scarce in our study domain (and in general, globally, Burrows et al. (2014)) and mainly found in costly planning units, not including representation targets for trajectory classes fails to ensure that they are represented.

It is challenging to predict the adaptive capacity of individual species to climate change, which depends on their historical experience and life-history characteristics (Morikawa & Palumbi, 2019). Instead, representing climate-velocity trajectory classes is a simple and easy-to-use method to meet this climate-smart objective at a community level (Brito-Morales et al., 2018), building ecological resilience (Mcleod et al., 2019) and improving the adaptive capacity of ecosystems under a spectrum of future ecological conditions (Roberts et al., 2017; Tittensor et al., 2019). They are also relatively straightforward to calculate now using the recent VoCC package in R (García Molinos et al., 2019).

### 3.3 Applicability and limitations

The use of climate velocity in spatial prioritization allows the incorporation of broadly applicable climate-change metrics into the design of networks of protected areas, along with other important considerations. However, caveats remain. For example, climate velocity is best suited for planning at the biogeographic scale or for identifying networks of large protected areas (Fredston-Hermann et al., 2018), and it is mainly applied in conservation to identify macro-refugia (Stralberg et al., 2018). There are challenges here, because our methods and case study highlight areas in the Mediterranean Sea that are climate refugia and that minimise future changes in species interactions, which can provide valuable information as initial large-scale prioritization to guide local conservation efforts. How this information translates for local conservation designs remains an open question. Nevertheless, as nations initiate negotiations for the Post-2020 Global Biodiversity Framework (CBD, 2020), climate-change mitigation and adaptation are increasingly important considerations. Moreover, in the ocean, most future gains in protected-area coverage will need to be in open-ocean waters, both within exclusive economic zones and beyond. This study provides a clear opportunity to incorporate climate velocity to directly address these needs at broad scales.

Importantly, our approach does not incorporate other aspects of climate change relevant for the design of networks of protected areas. Climate connectivity, for example, is important because it quantifies paths that facilitate species’ movement to suitable future climates. Climate-velocity trajectories could be used to create a climate-connectivity matrix and prioritize climate connections following a similar approach to that developed to account for larval dispersal (Beger et al., 2010). Also, because we know that climate velocity magnitudes and directions differ through the water column (Brito-Morales et al., 2020) this could be captured if protected-area policy allowed for three-dimensional zoning in the ocean (Brito-Morales et al., 2018; Venegas-Li et al., 2018). Finally, because slow-moving areas are not necessarily regions with less impact due to slow warming rates (i.e., areas where climate warming changes less) or less prone to catastrophic events (e.g., heatwaves), our approach does not incorporate a measure of impact (see supporting information).

Our approaches outlined here provide a framework for climate-smart conservation planning that complements conventional conservation-planning priorities, including the representation of biodiversity and the minimization of associated costs. Importantly our methods depend exclusively on widely accessible data and free-to-use, open-source software, making them available for use in almost any situation and encouraging other research groups to advance the field, with the aim of delivering truly robust networks of climate-smart protected areas by 2030.

## ACKNOWLEDGEMENTS

N.A.D. is supported by the Fundación Bancaria ‘la Caixa’ Postgraduate Fellowship (LCF/BQ/AA16/11580053) and by the University of Queensland Research Training Scholarship. I.B.M. is supported by the Advanced Human Capital Program of the Chilean National Research and Development Agency (grant no. 72170231). We are deeply thankful to Tessa Mazor for sharing the opportunity cost layer for the Mediterranean Sea, and to AquaMaps team for providing the most updated species distribution data. The authors declare that there is no conflict of interest.

## AUTHOR CONTRIBUTIONS

N.A.D and I.B.M conceived and designed the research with inputs from D.S, H.P, C.K, and A.R. I.B.M and N.A.D analyzed the data and D.S, H.P, and A.R contributed in the discussion of ideas and analyzes. N.A.D and I.B.M wrote the first draft. All authors contributed and commented on the manuscript.

## DATA AND CODE AVAILABILITY

Data and R scripts are available at Zenodo under the identifier http://doi.org/10.5281/zenodo.3875796.

