## Supporting Information for "Incorporating climate velocity into the design of climate-smart networks of protected areas"

### SUPPLEMENTARY MATERIALS & METHODS

#### Opportunity costs

The combined opportunity cost data used for our analyzes represents commercial, recreational, and aquaculture sectors for the Mediterranean Sea (Mazor et al., 2014). Commercial fishing data are based on fish tonnage caught over 28 geographical regions provided by the General Fisheries Commission for the Mediterranean and the average market value of species. Non-commercial fishing data are based on the number of recreational fishers per country, the cost of expenditure on fishing gear, and the value of catch per year within 12-nautical-mile territorial waters. Aquaculture cost estimates are based on the production multiplied by the market value of two primary aquaculture species in the Mediterranean Sea.

#### Calibrating Marxan parameters

Marxan provides alternative solutions that minimize the total score of the Marxan objective function, which is given by the sum of the total cost of the selected planning units, a penalty assigned for features that do not meet their targets, and a score related to the perimeter of the spatial configuration of the reserve system. Parameters we explored in this calibration analysis are the number of iterations, the species penalty factor (SPF), and the boundary-length modifier (BLM). The number of iterations determines how close Marxan gets to the optimal solution. The SPF is a parameter that forces conservation features to meet their targets. The BLM determines the level of aggregation across planning-unit boundaries in the solutions. To ensure a consistent and reasonable level of fragmentation for the spatial configuration of Marxan solutions that allows comparison among scenarios, we developed a tailored calibration and selection process:

**Step 1:** For the initial exploratory analysis, we used the Base scenario to calibrate the number of iterations. We chose a range of 10 different BLM values to run 10 scenarios. To ensure that all targets were being met in each scenario, we increasd the SPF parameter until this was achieved. We ran each scenario for three iteration values: $10^6$, $10^7$, and $10^8$, and for 10 times instead of 100, for speed purposes.

**Step 2:** We plotted the cost vs. fragmentation of the solutions to understand the relationship between these factors and the BLM values (Figure S1).

$$Fragmentation = \frac{Perimeter\ of\ the\ solution}{Perimerter\ of\ the\ solution\ if\ it\ were\ a\ circle}$$

By increasing the number of iterations, the solutions become more similar as they get closer to optimality (decreases variability and the cost, perimeter, and fragmentation of the solutions) (Figure S1). Beyond some point, the extra time required by a higher number of iterations will yield no substantial improvement in the solutions. The higher the number of iterations is, the longer the scenarios will take to run, so we recommend using a low number of runs for this calibration phase, such as 10 for each scenario.

Based on this analysis we choose an iteration value of $10^7$ as it substantially improves the optimality of the solutions compared to $10^6$ iterations. The computational time to run the scenarios with 100 runs with $10^7$ iterations is several hours. If we increase to $10^8$ iterations, minor improvement of the solutions comes with an increase in computational time to days.

**Step 3**: We run 10 BLM scenarios for each approach explored in this study (Table 1) to choose a similar level of fragmentation for all scenarios. We ensured that all representation targets were being met by increasing the SPF, when needed. We found that the optimal BLM values were able to accomplish a fragmentation value between 5 and 6, before substantially increasing the cost of the solutions. Therefore, we chose those BLM values for each approach where the ten solutions had fragmentation values ranging between 5 and 6.

**Calibrating Velocity as a Feature scenario**

To find the optimal trade-off between cost, fragmentation, and area of the solution, we calibrated the representation target assigned to the slowest-moving planning units for Velocity as a Feature scenario. We ran 11 scenarios by increasing the representation target for slow-moving planning feature from 0 to 100%. For each of the 11 scenarios, we first calibrated the boundary-length modifier as in Step 3, to ensure a fragmentation value of the solutions between 5 and 6. After we calibrated the BLM, we ran each scenario 10 times. On this basis, we chose to represent 30% of slow-moving feature.

### SUPPLEMENTARY DISCUSSION

#### Choosing planning unit shape and size

In our analysis, we used 0.25º grid square cells as our planning units shape because: 1) this is the finest resolution of global climate models; 2) bigger resolutions such as 0.5º could be too coarse for some protected-area planning; and 3) they match the grid shape of climate models. However, the shape of a planning unit can vary, and the selection of different shapes might have implications when using climate velocity as a boundary in Marxan (e.g., Velocity as a Boundary scenario). For example, in the marine realm, multi-edged planning units (e.g., hexagons) might be more efficient than a square grid because they better represent the land-sea transition in coastal regions as sites have six sides shared among individual units rather than four edges. As climate velocity is usually calculated in a regular grid square, the use of a hexagonal planning-unit grid would create two extra boundaries that were not originally considered in the estimation of local climate velocity. In this case, and for other irregular planning-unit shapes, the simplest solution is to use the same climate-velocity magnitude for the extra sides. These aspects were beyond the scope of the current method but could be considered in further research.

**Extended applicability and limitations**

Climate refugia defined on the basis of climate velocity are not necessarily regions with less impact due to slow warming rates or extreme events. This is a fundamental distinction because climate velocity is calculated as the ratio of the temporal trend over the corresponding spatial gradient in climate. In this context, fast warming rates (i.e., steep temporal trends) can be irrelevant in areas with steep spatial gradients (i.e., more heterogeneous thermal landscapes), which can lead to slow climate velocities (Brito-Morales et al., 2018). For instance, considering two planning units, one with a slow warming rate (0.5°C per unit time) and gentle spatial gradient (10°C $km^{-1}$) and the other with fast warming rate (1°C per unit time) and steep spatial gradient (100°C $km^{-1}$). In this case, the second planning unit has twice the warming rate of the first one but a slower climate velocity due to the steep spatial gradient experienced in that area.

Another important consideration is the compatibility of our approach with other aspects of protected-area design, such as ecological connectivity among protected areas. For example, it is not possible to simultaneously use Velocity as a Boundary scenario with the approach developed to account for asymmetric connectivity (Beger et al., 2010), as both methods modify the BLM. However, Marxan connect (Daigle et al., 2020), a recently developed tool allows incorporation of ecological connectivity as a feature, which is compatible with our Velocity as a Boundary approach. Nevertheless, we can simultaneously use the approach of Beger et al. (2010) with our Velocity as a Feature, as this scenario is independent of the BLM. Further analysis could determine which of these two options are best at selecting slower-moving areas while maximizing ecological connectivity among planning units.

**SUPPLEMENTARY FIGURES AND TABLES**

TABLE S1 List of phyla included in the Marxan analysis at the Mediterranean with their corresponding number of species computed using a threshold of 0.5 probability of occurrence.

| Phylum | Class | Total |
|---|---|---|
| Annelida | Polychaeta | 169 |
| Arthropoda | Branchiopoda | 2 |
| | Malacostraca | 114 |
| | Maxillopoda | 30 |
| | Ostracoda | 2 |
| | Pycnogonida | 12 |
| Brachiopoda | Articulata | 4 |
| | Inarticulata | 1 |
| Bryozoa | Gymnolaemata | 53 |
| Chaetognatha | - | 1 |
| Chlorophyta | Bryopsidophyceae | 8 |
| | Ulvophyceae | 12 |
| Chordata | Actinopterygii | 342 |
| | Appendicularia | 6 |
| | Ascidiacea | 45 |
| | Cephalochordata | 1 |
| | Elasmobranchii | 50 |
| | Mammalia | 2 |
| | Reptilia | 12 |
| | Thaliacea | 7 |
| Cnidaria | Anthozoa | 68 |
| | Cubozoa | 1 |
| | Hydrozoa | 73 |
| | Scyphozoa | 6 |
| Ctenophora | Tentaculata | 1 |
| Cyanobacteria | - | 2 |
| Echinodermata | Asteroidea | 12 |
| | Crinoidea | 1 |
| | Echinoidea | 10 |
| | Holothuroidea | 11 |
| | Ophiuroidea | 9 |
| Echiura | - | 1 |
| Foraminifera | Polythalamea | 1 |
| Gastrotricha | - | 18 |
| Gnathostomulida | - | 2 |

| | | |
|---|---|---|
| Hemichordata | Enteropneusta | 1 |
| Mollusca | Aplacophora | 1 |
| | Bivalvia | 170 |
| | Cephalopoda | 16 |
| | Gastropoda | 98 |
| | Polyplacophora | 4 |
| | Scaphopoda | 5 |
| Ochrophyta | Phaeophyceae | 8 |
| Phoronida | - | 3 |
| Platyhelminthes | Turbellaria | 3 |
| Porifera | Calcarea | 8 |
| | Demospongiae | 30 |
| Rhodophyta | Compsopogonophyceae | 1 |
| | Florideophyceae | 25 |
| | Rhodellophyceae | 1 |
| Sipuncula | Phascolosomatidea | 7 |
| | Sipunculidea | 1 |

TABLE S2 Final SPF and BLM values used to run Marxan scenarios for each approach.

| Scenarios | SPF | BLM |
|---|---|---|
| 1. Base | 1.8 | 250 |
| 2. Velocity as a Cost | 1.5 | 0.0001 |
| 3. Velocity as a Boundary | 1.6 | 7500000 |
| 4. Velocity as a Feature | 1.8 | 250 |
| 5. Trajectories as a Feature | 1.8 | 250 |

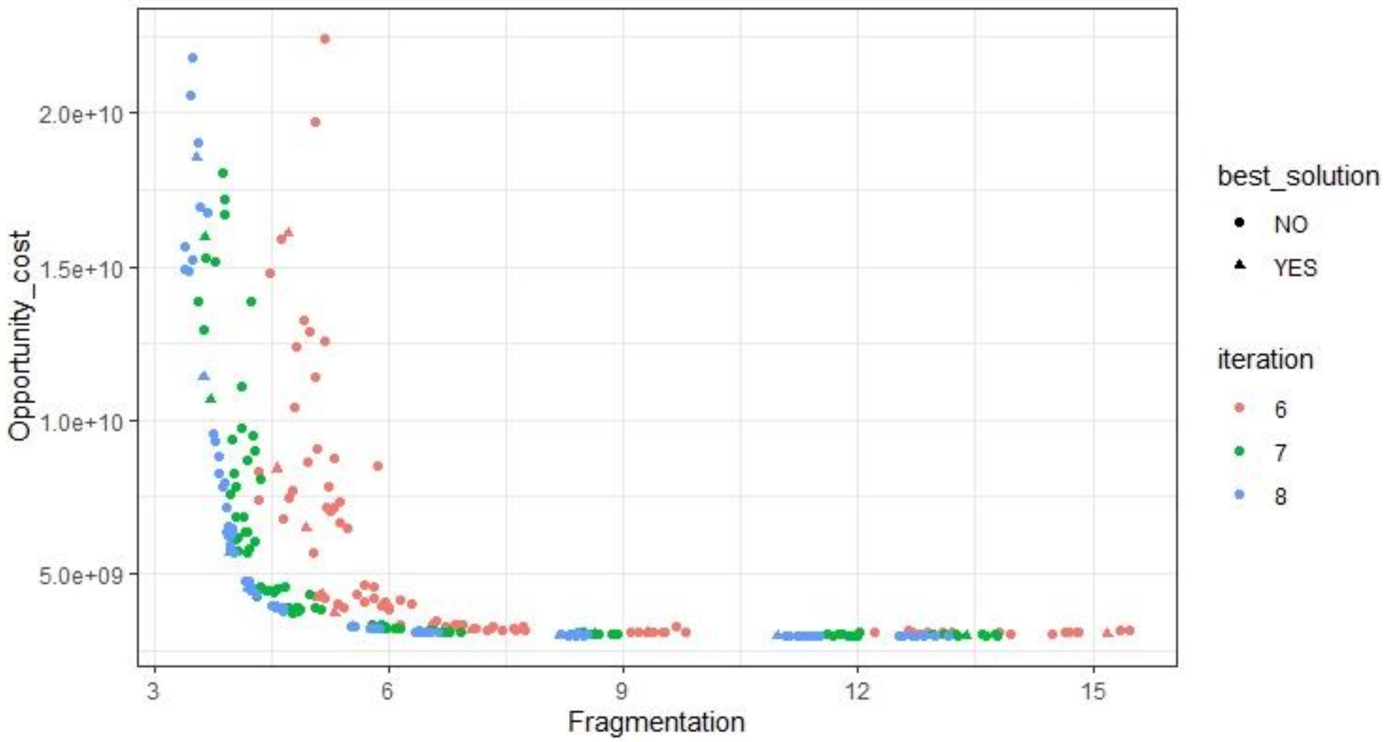


FIGURE S1 Trade-off curve for the cost and fragmentation, for the Base scenario, for three different iteration values of a set of 10 BLM Marxan scenarios.

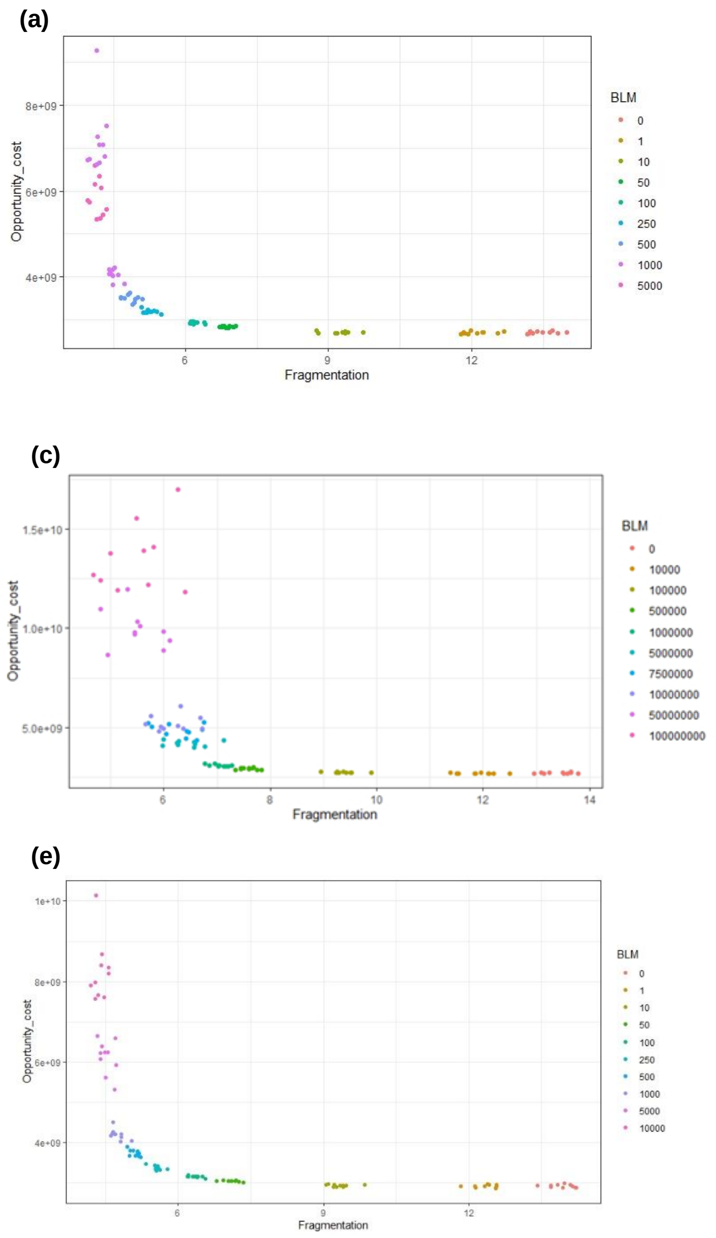


FIGURE S2 Trade-off curve for the cost and fragmentation for (a) Base, (b) Velocity as a Cost, (c) Velocity as a Boundary, (d) Velocity as a Feature, and (e) Trajectories as a Feature scenarios, for a set of 10 BLM Marxan scenarios.

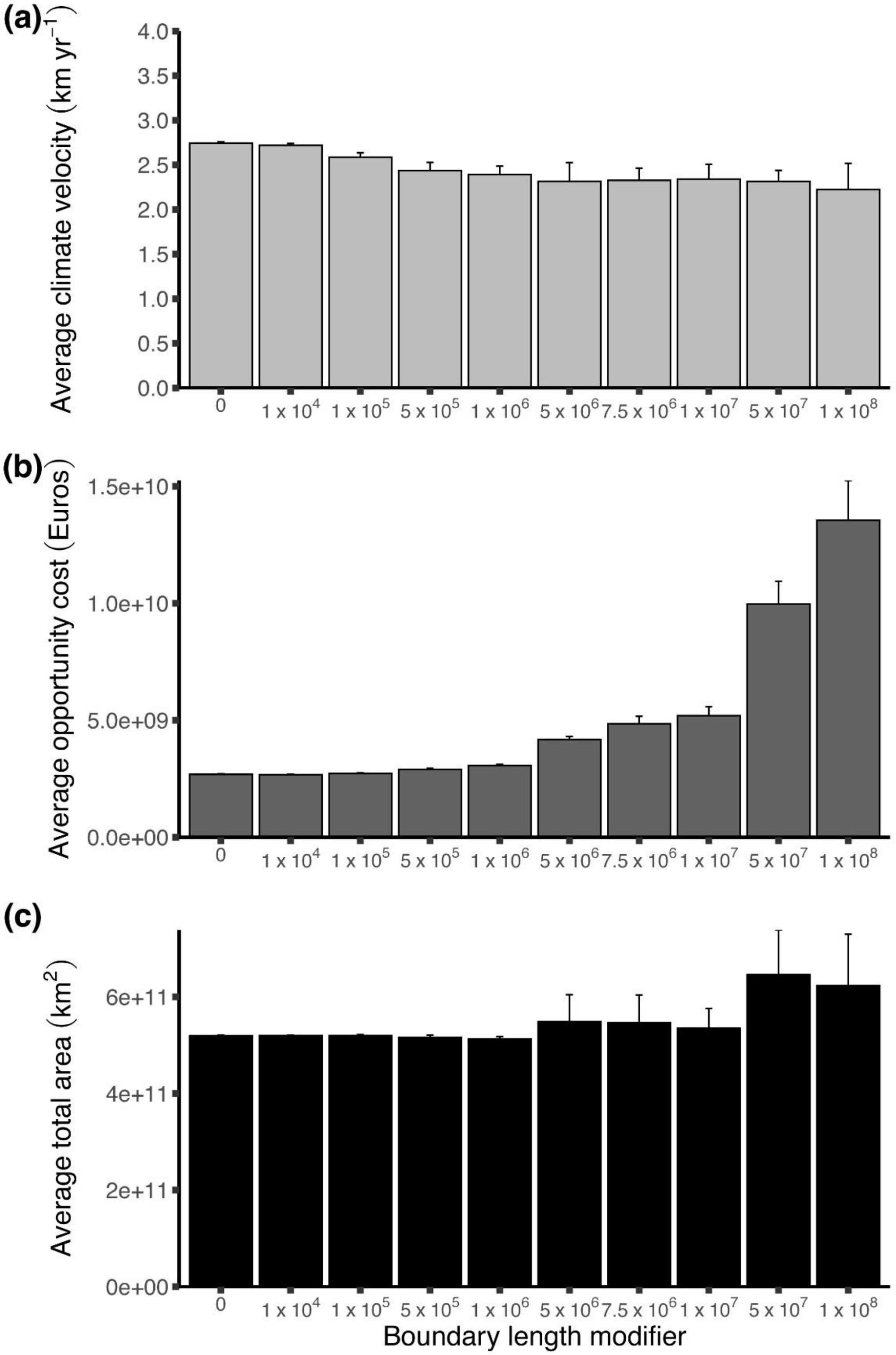


FIGURE S3 **(**a) Median climate velocity, (b) opportunity cost in monetary value (n = 10 ± s.d) and (c) total area for each BLM for Velocity as a Boundary scenario for each of Marxan 10 solutions.

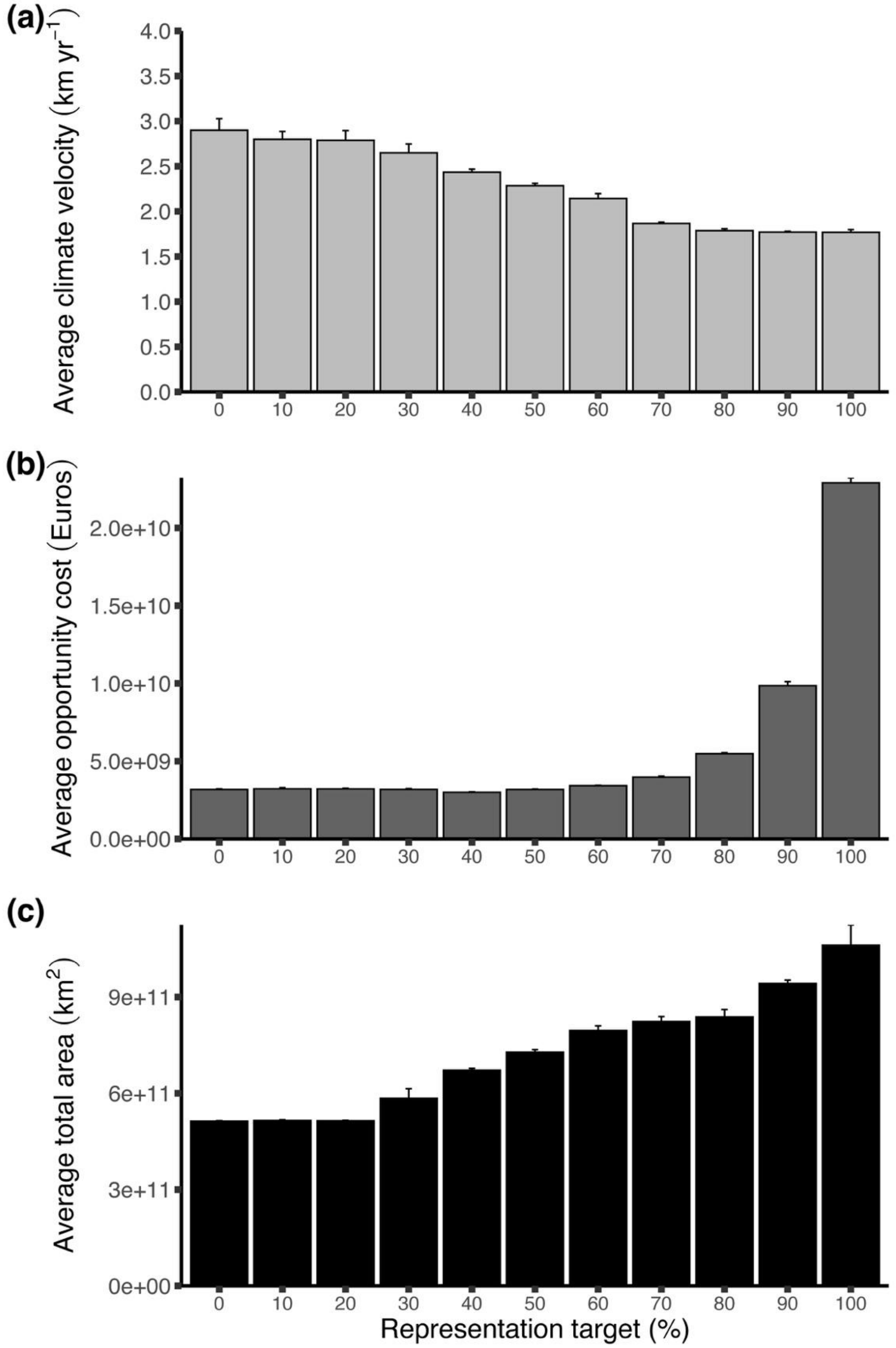


FIGURE S3 **(**a) Median climate velocity, B) opportunity cost in monetary value (n = 10 ± s.d) and C) total area for each representation target Velocity as a Feature scenario for each of Marxan 10 solutions.
